# Iliac vein morphology and wall shear stress: a statistical shape modelling and CFD analysis of patient-specific geometries

**DOI:** 10.64898/2026.02.17.706277

**Authors:** Magdalena Otta, Karol Zając, Ian Halliday, Chung S. Lim, Maciej Malawski, Andrew J. Narracott

## Abstract

Deep vein thrombosis (DVT) is a prevalent vascular condition in which venous anatomy and flow disturbances contribute to the risk of thrombosis, but the mechanistic links between vessel shape and haemodynamics remain poorly quantified. Although computational fluid dynamics (CFD) can estimate flow-related risk metrics such as low wall shear stress (WSS), the influence of anatomical fidelity on these predictions is not well understood. Statistical shape modelling (SSM) offers a principled framework for characterising geometric variability, but its integration with CFD in venous applications is still emerging. This study investigates how different levels of anatomical representation—2D projections, simplified 3D extrusions, and full 3D reconstructions of the common iliac veins—influence both the statistical structure of venous shape variability and the haemodynamic metrics derived from CFD. Using patient-specific MRI/CT data from twelve cases, we constructed SSMs in Deformetrica and performed steady-state CFD simulations in ANSYS Fluent under standardised inflow conditions. We compared the variance structure of the 2D and 3D latent spaces and quantified correlations between principal shape modes and low-WSS burden across three thresholds (≤ 0.05, 0.10, 0.15 [Pa]). Idealised 3D geometries consistently produced larger low-WSS areas than patient-specific shapes, with average increases of 118–136% across thresholds. The 2D SSM exhibited a strongly hierarchical variance spectrum with one dominant mode that correlated significantly with WSS, whereas the 3D SSM showed a flatter spectrum with weaker univariate associations. These findings demonstrate that geometric fidelity and alignment strategy critically influence shape–flow relationships, highlighting the need for careful model selection when using CFD-based haemodynamic indicators in DVT research.

**Author summary:** Deep vein thrombosis (DVT) is a common condition in which blood clots form in the deep veins of the leg and can lead to serious long-term complications. Although medical imaging captures important anatomical differences between patients, it remains unclear how these variations in vein shape influence local blood flow and the associated risk of clot formation. To address this challenge, we developed a computational framework that combines statistical shape modelling (SSM) with computational fluid dynamics (CFD) to analyse the relationship between venous geometry and haemodynamic risk factors.

We examined the common iliac veins at three levels of anatomical detail: simplified two-dimensional projections, intermediate three-dimensional extrusions, and full three-dimensional reconstructions derived from MRI/CT data. By comparing these representations, we show that geometric fidelity strongly affects both the detected modes of anatomical variation and the resulting flow predictions. Simplified geometries consistently overestimated regions of low wall shear stress, a flow feature associated with thrombosis, compared to full 3D models. We also found that shape–flow associations depend heavily on how shapes are aligned and represented.

Our findings highlight the importance of anatomical detail in computational venous modelling and provide a foundation for more personalised, simulation-based tools to support DVT treatment.

## Introduction

Deep vein thrombosis is a common disease that affects approximately 1 in 20 individuals in the general population. Among those diagnosed, approximately one third experience recurrent episodes and 20-50% develop post-thrombotic syndrome (PTS). Iliofemoral DVT is typically associated with the most severe symptoms, including swelling and pain in the legs, due to limited collateral venous drainage in the region. This form of DVT also has the highest risk of developing PTS [1]. Alterations in blood flow represent one of the three main risk factors for thrombosis, but remain the least understood factor. Haemodynamics is known to influence thrombus initiation and progression with low wall shear stress (WSS) related to blood coagulation and clot formation [2]. In addition, the efficacy of invasive interventions, such as stenting, relies on adequate inflow to maintain patency. Venous anatomy varies between individuals, but can also differ between the left and right limbs of the same person [3]. It is not clear to what extent these differences influence the local and systemic characteristics of blood flow that promote the development of thrombosis.

Despite its relevance, haemodynamics is currently not quantitatively assessed in routine care. Treatment decisions in DVT are largely based on anatomical images from MRI/CT, ultrasound, and angiography. Flow characteristics are not measured but could be estimated with simulation. Computational fluid dynamics (CFD) and statistical shape modelling (SSM) have the potential to improve the understanding of links between venous anatomy and associated changes in haemodynamics. SSM alone enables the quantification of complex anatomical configurations and supports the examination of structural variability. A combined framework that integrates CFD-derived metrics, such as vessel regions exposed to low wall shear stress, with SSM-derived shape descriptors could support haemodynamic-based risk stratification for patients treated for deep vein thrombosis (DVT) as previously proposed [4].

Previous studies have explored the use of CFD to simulate venous flow and assess thrombosis risk, including patient-specific analyses of iliac vein compression and stenosis-severity effects [5, 6], while SSM has been applied to characterise anatomical variability in various vascular contexts. Bruse et al. [7] focused on linking the 3D anatomical shape characteristics of the repaired aortic arches to functional cardiac metrics, such as the ejection fraction and blood pressure gradients. They used SSM to extract shape biomarkers and then correlated them with clinical haemodynamic measurements obtained from imaging and patient data, but did not perform CFD simulations. Pajaziti et al. [8] introduced a deep learning approach that predicts 3D pressure and velocity fields in the aorta using shape features derived from statistical shape modelling, trained on 3,000 CFD simulations of synthetic shapes. Their framework replaces traditional CFD with a fast surrogate model. Recent work by Mazzoli et al. [9] combined SSM and CFD to study how variability in aortic shape affects haemodynamics, identifying correlations between shape modes and metrics such as wall shear stress. Together, these studies demonstrate the value of shape-informed flow analysis in arterial disease. Our work extends this concept to venous anatomy, focussing on DVT and systematically varying anatomical fidelity—how closely the geometry used in the analysis preserves the true, patient-specific anatomy—to evaluate flow metrics related to thrombosis, addressing a critical gap in venous haemodynamics research.

Deformetrica [10] is a dedicated Python library commonly used for SSM analysis. The tool is built upon the mathematical framework of large-deformation diffeomorphic metric mapping (LDDMM), which models anatomical shape variability through smooth, invertible transformations known as diffeomorphisms [11]. In the LDDMM framework, shapes are treated as points on a Riemannian manifold, with deformations represented by geodesic paths, the shortest routes, connecting them. This formulation enables robust and consistent comparison of shapes, supporting statistical analysis of anatomical variability. Deformetrica implements models that include atlas construction and geodesic-based dimensionality reduction [12]. Two are of particular interest in this research, the Deterministic Atlas and Principal Geodesic Analysis. The deterministic atlas framework estimates a representative template shape that minimises the sum of squared geodesic distances to all observed shapes within a population. Each individual shape is modelled as a smooth deformation of this template, governed by initial momenta that encode both the direction and magnitude of variation. Optimisation balances the fidelity to the data with the regularity of the deformation, yielding a mean anatomical shape and a set of geodesic paths that capture individual variability across the cohort. Once the atlas is constructed, Principal Geodesic Analysis (PGA) can be applied to the resulting deformation momenta to identify dominant modes of shape variation. These principal geodesics offer a compact and interpretable representation of anatomical variability that can be further correlated with clinical or haemodynamic metrics.

In the context of blood flow modelling for DVT, these SSM techniques can help quantify vascular deformation patterns and support simulation-based treatment strategies. This study builds on previous work [4], which examined the venous anatomy using two-dimensional SSM integrated with CFD. In the present work, we advance our previous 2D shape–flow analysis by applying statistical shape modelling to full three–dimensional iliac vein geometries. These 3D shapes correspond directly to the 2D projections analysed in the earlier study, and we additionally consider 3D geometries that retain their native out–of–plane curvature extracted from MRI/CT data. These geometries represent three levels of anatomical fidelity; 2D projection, simplified 3D extrusion, and full 3D reconstruction.

The central hypothesis of this study is that the relationship between venous morphology and haemodynamics depends strongly on the level of geometric fidelity used to represent the anatomy. Specifically, we hypothesise that simplified or projected geometries capture only a subset of the morphological variation that influences wall shear stress, whereas anatomically richer three–dimensional models may reveal more complex shape–flow associations. To evaluate this, we address three research questions: (1) How does geometric fidelity influence the statistical structure of venous shape variation obtained through SSM? (2) How do these differences in shape representation affect the association between principal shape modes and regions of low wall shear stress derived from CFD? (3) Are particular anatomical deformation patterns consistently linked to haemodynamic risk indicators across fidelity levels? Addressing these questions is informative for future development of clinical workflows to assess DVT patients, with significant implications for the choice of imaging protocol used.

By systematically varying anatomical representation and analysing the resulting shape–flow relationships, this study aims to clarify how anatomical details affect haemodynamic metrics in the context of deep vein thrombosis. Beyond demonstrating a combined SSM–CFD workflow for venous anatomy, we explicitly quantify how model fidelity and alignment reshape the latent variance structure and the strength/localisation of shape–flow associations—an aspect not previously measured in venous modelling.

## Materials and methods

Our methodology follows the workflow presented in Fig. 1. The clinical data used in this study consisted of 3D volumetric MRI and CT scans from patients treated for DVT. From these images, we reconstructed the 3D geometry of the common iliac vein and generated their 2D projections in the plane corresponding to standard angiographic imaging used in clinical practice. For selected parts of the analysis, we also created simplified 3D extrusions based on the 2D projections, neglecting out-of-plane curvature and assuming that the centreline radii measured in the 2D projections accurately represent the corresponding 3D radii.

**Fig 1.**
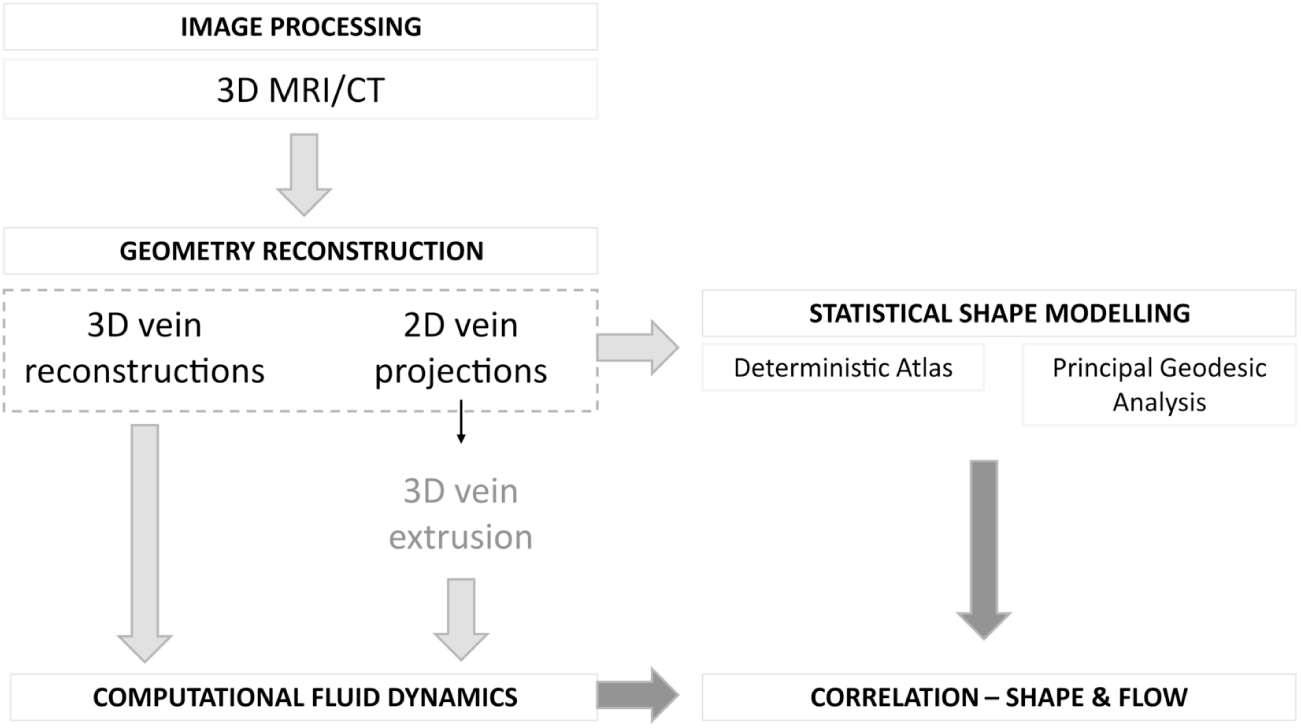
Workflow overview. Methodology applied from segmentation of medical images, through 2D and 3D shape construction, to statistical shape analysis and 3D CFD simulations.

Two complementary modelling approaches were used: statistical shape modelling (SSM) and computational fluid dynamics (CFD). We compared results at different levels of geometric complexity and assessed statistical correlations to investigate possible relationships between venous shape and flow-related metrics, specifically wall shear stress.

Medical images were processed with custom Python tools to reconstruct patient-specific venous geometries - CFD simulations were conducted using ANSYS Fluent 2024 R2, and statistical shape modelling (SSM) was performed with Deformetrica. Details of each step are provided in the following sections.

### Image processing

A set of 12 3D geometries of the common iliac veins was extracted from volumetric MRI and CT images by drawing contours of the vein lumens along the length of the vein. The unification of the internal and external iliac veins and a merging of the common left and right iliac veins, into the inferior vena cava (IVC), were used to ensure the same anatomical scope of the vessel of interest. A custom Python script was written to draw and save the contours (Fig. 2).

**Fig 2.**
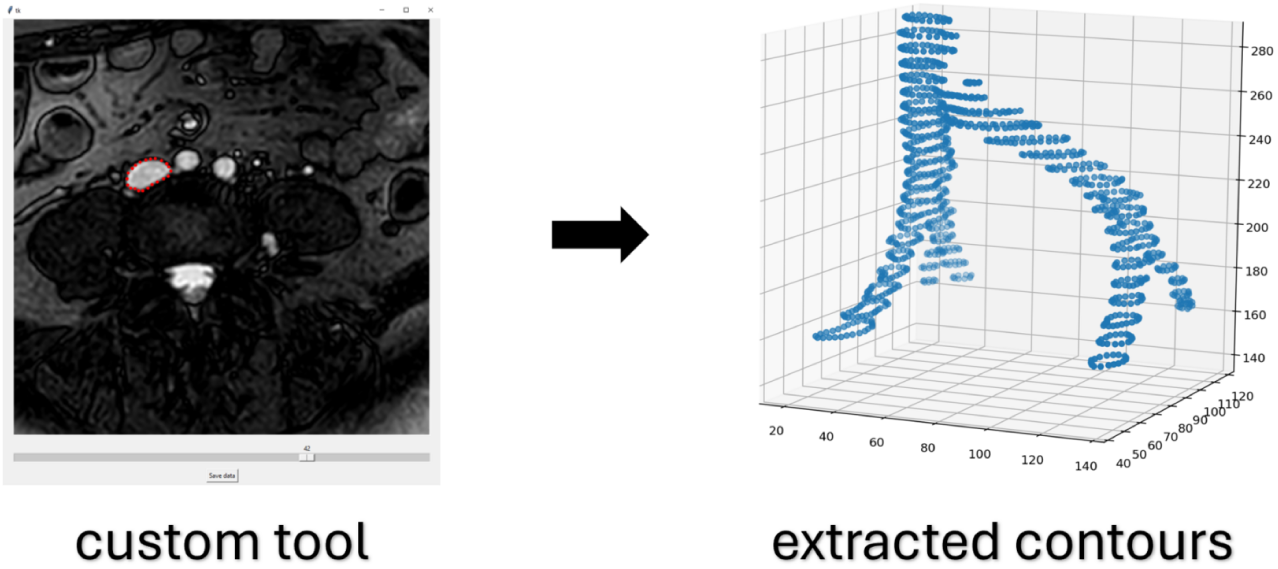
Extraction of venous contours. Contour of venous lumens extracted from 3D volumetric images using custom Python tool.

### Geometry reconstruction

Three types of geometries (Fig. 3) were constructed based on the information available from medical images:

A. full 3D shapes of common iliac veins from MRI/CT images,
B. 2D contours depicting the projection of the 3D geometries into the plane of the angiogram plane,
C. 3D extrusions of the 2D projections (These simulate a workflow for 3D reconstruction, based solely on 2D angiograms.)

**Fig 3.**
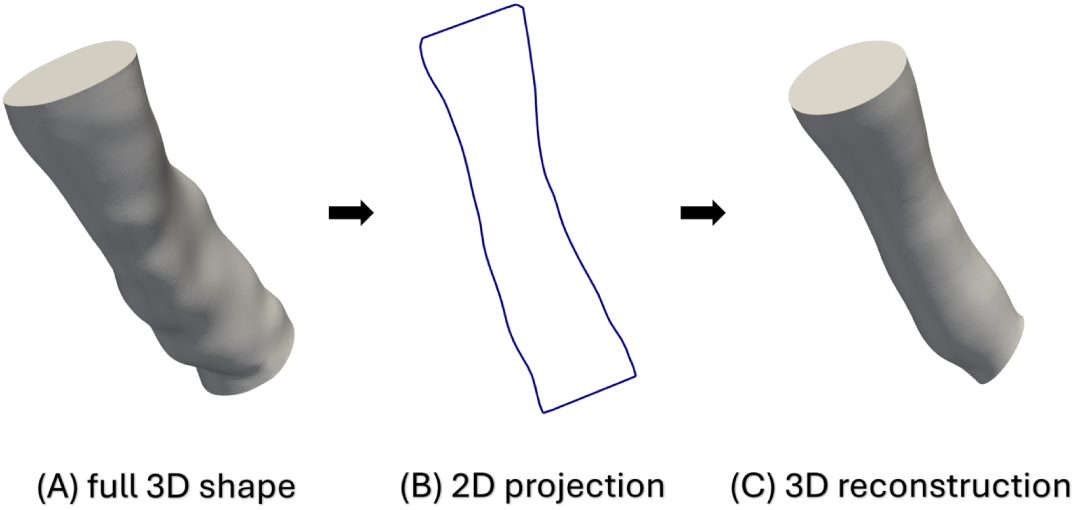
Shape types. (A) full 3D shape from volumetric images; (B) 2D projection in angiogram plane; (C) 3D extrusion of 2D shape.

Our aim was to compare how much plausible (in the context of a clinical workflow) geometric information - effectively, what type of imaging - is sufficient to link shape modes to haemodynamic parameters computed in 3D CFD simulations.

The 3D geometries in (A) were built in STL format, based on the extracted contours using a custom Python tool. They were later processed for simulations in ANSYS SpaceClaim 2024 R2, including smoothing and regularisation, and converted into VTK format for SSM. The 2D projections in (B) were extracted from the 3D VTK files and interpolated to a consistent number of points and orientation. A custom script was used to reconstruct the simplified 3D geometries in (C), based on the 2D projections in STL format. They were later processed for simulation following the workflow from (A).

### Computational fluid dynamics

All 3D venous geometries were analysed using an ANSYS framework, covering all stages from geometry preparation in SpaceClaim 2024 R2, through meshing and simulation in Fluent 2024 R2, to the output analysis in CFD-Post 2024 R2.

A polyhedral mesh was created for each 3D shape in set (A) and (C) with an element size of 0.20 *mm* (average vein diameter ∼ 13*mm*), determined by mesh sensitivity analysis. A mesh-refinement study showed that low-WSS areas differed by less than 1% between the two finest meshes (0.25 − 0.20 *mm*) across all thresholds, indicating practical mesh independence at a characteristic element size of 0.20 mm (see Supplementary material Fig. S1 and Tab. S1).

Simulations were performed in the steady-state, assuming laminar flow with constant zero pressure set at the outlet and a parabolic velocity profile at the inlet, estimated from previous work [4] based upon a 0D lumped-parameter model - baseline 0.0782*m/s*. A variation in the inlet velocity ±20% was applied uniformly across all shapes to assess sensitivity to inflow conditions. Additionally, the mean *Re*, equal to 354, of the baseline velocity simulations was used for a further comparative analysis between geometries. The blood constitutive model corresponded to incompressible Newtonian fluid of constant density *ρ* = 1050*kg* · *m*^−3^ and viscosity *µ* = 0.0035*Pa* · *s*.

The CFD workflow was automated using the PyFluent library, including batch processing for the meshing and simulation steps.

The metric of interest was the area of the vein wall subject to low wall shear stress bounded by three thresholds (≤ 0.05, 0.10, 0.15[*Pa*]), to which the data were filtered. An example of such a banded region is shown in Fig. 4. Clinical thresholds for venous WSS have not been standardised; therefore, we analysed three nested low-WSS cut-offs spanning the range associated with low-shear venous coagulation in the literature. These thresholds should be interpreted as windows on the same underlying low-shear distribution, not as clinically validated cut-offs.

**Fig 4.**
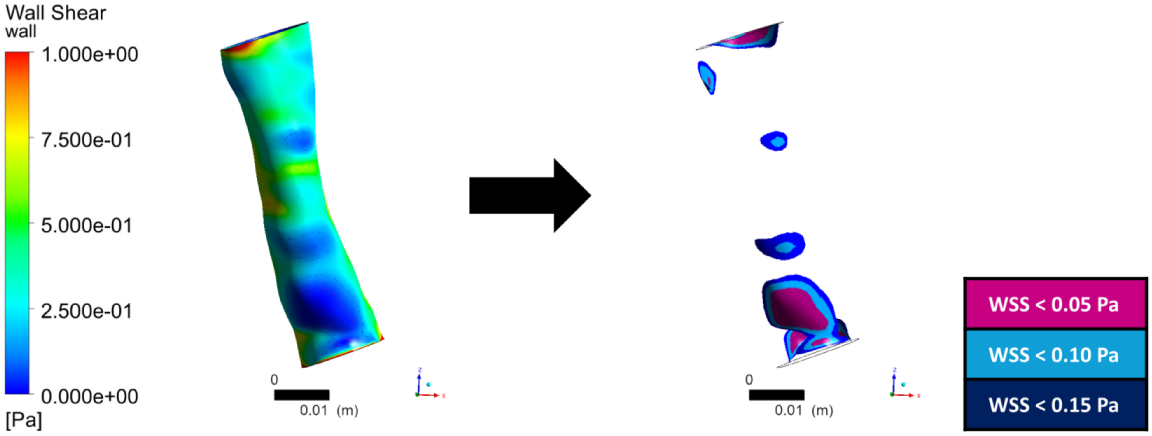
An example of wall shear stress distribution (left) with bands of WSS below three thresholds filtered out (right).

To confirm that steady-state low-WSS regions captured the same haemodynamic structures as oscillatory metrics, we performed a transient pulsatile simulation using a sinusoidal inflow waveform (*f* ≈ 1.33 *Hz*). Regions of elevated OSI coincided spatially with the steady-flow low-WSS areas, indicating that transient effects did not alter the organisation of low-shear zones (see Supplementary material Fig. S2).

### High-Performance Computing workflow and infrastructure

To eliminate interactive graphical user interface operations and minimise manual intervention, the entire simulation pipeline was implemented using the PyFluent Python library. All simulations were performed using a fully automated, script-based workflow designed to ensure scalable and efficient utilisation of resources. This approach allowed programmatic definition, parametrisation, and execution of ANSYS Fluent workflows. A CSV file with simulation-specific parameters governing meshing and solver setup enabled systematic and reproducible execution of simulation scenarios across all geometries considered.

Computational resources were provided by the Ares High-Performance Computing (HPC) cluster at ACC Cyfronet (Kraków, Poland). The ANSYS Fluent module installed on the cluster supports distributed-memory parallelism, and Intel MPI was used to accelerate each individual simulation. Although MPI parallelism was employed at the single-simulation level, all simulations were independent of one another. Therefore, an array job strategy available in the SLURM queuing system was adopted. The strategy of submitting as a separate task with a fixed and relatively small resource allocation improved overall cluster throughput, reducing queue waiting times, and enabling straightforward monitoring or recovery of individual runs. Each simulation was executed using 8 CPU cores and 32 GB of RAM, resulting in an average combined meshing and solver execution time of approximately 15-20 minutes per case.

A single batch submission ensured consistency across all cases and the configuration provided an effective balance between computational efficiency and resource usage for this study. The simulation results and the meshes were stored in long-term storage in HDF5-based formats for efficient data handling. Defining quantities for post-processing and exporting solver results directly within PyFluent ensured automated and consistent extraction of relevant haemodynamic metrics for analysis.

### Statistical shape modelling

Statistical shape modelling was performed using the Deterministic Atlas (DA) and Principal Geodesic Analysis (PGA) framework implemented in Deformetrica. In this approach, the DA step is essential, as it estimates a population template *I*_0_ and the initial momenta 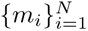 that deform this template to each subject in the cohort. These momenta provide a common reference representation of deformation; without such a template–centred framework, the shapes would remain in incompatible coordinate systems and no meaningful statistical analysis could be conducted. The atlas seeks a template and set of deformations that minimise an energy functional balancing data fidelity and deformation regularity:

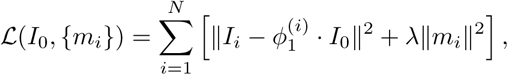

where *I_i_* denotes the observed shape of subject *i*, 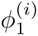 is the diffeomorphic flow generated by *m_i_*, and *λ* controls the regularity of the deformation. This optimisation yields a mean anatomical shape together with geodesic paths that describe the deformation from the template to each subject, following the Large Deformation Diffeomorphic Metric Mapping (LDDMM) framework of Beg et al. (2005) [11].

PGA generalises principal component analysis to the non-linear manifold of diffeomorphic deformations by analysing variability in the tangent space at the template [13]. The covariance of the initial momenta,

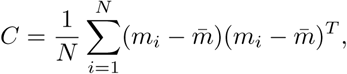

is decomposed to obtain the principal modes of shape variation, each associated with an eigenvalue *λ_k_* and eigenvector defining a geodesic direction of deformation (see also Durrleman et al. 2014 [12] for a Deformetrica implementation). For each mode *k* and subject *i*, Deformetrica outputs a raw PGA coefficient *c_ik_*, which is expressed in arbitrary tangent-space units. To facilitate interpretation, and to enable comparison across modes, these coefficients were converted into standard-deviation coordinates using

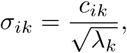

so that displacements of ±1*σ* correspond to one geodesic standard deviation along mode *k*. This parameterisation is used for visualising and quantifying the principal anatomical variations captured by the model.

The Deformetrica parameter configurations used for the 2D and 3D are provided in Supplementary material - Tables S2 and S3 respectively.

### Shape pre-processing for SSM

All 2D and 3D venous geometries were exported in legacy VTK format to ensure compatibility with Deformetrica, with a target mesh resolution of approximately 1 mm. For the 3D shapes, the vessel ends were first re-capped using planes parallel to one another, to provide a consistent truncation for ensuing statistical analysis; this modification was applied solely for SSM and did not affect the CFD-specific capping, which was orthogonal to the local vessel axes. In the 2D dataset, the anatomical orientation of the vessels was deliberately preserved: the shapes were aligned by translating them such that the inlet centre points lay along the global *x*-axis, thereby retaining the natural inflow angle and curvature. In contrast, the 3D shapes were aligned by minimising the pairwise discrepancy between their centrelines, a process that effectively removes global anatomical pose and isolates only the intrinsic geometric differences in local vessel morphology. This difference in alignment strategy reflects a practical limitation: attempts to apply the anatomically anchored 2D alignment approach to the 3D dataset led to non-convergent or unstable SSM atlas solutions in Deformetrica, necessitating the use of a more robust centreline-based method in 3D. The resulting alignments for both 2D and 3D datasets are shown in Fig. 5.

**Fig 5.**
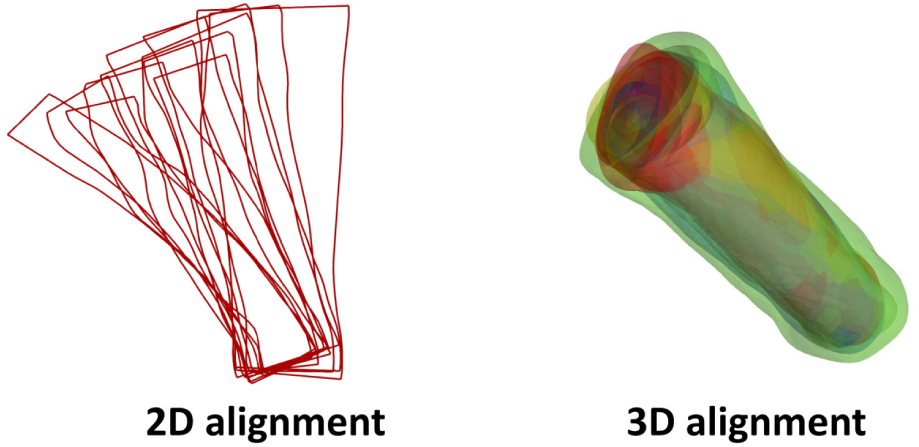
Alignment procedures applied to the 2D and 3D venous geometries prior to statistical shape modelling. Left: 2D shapes aligned by translating the inlet centre points onto a common horizontal axis, preserving the anatomical inflow orientation and overall pose. Right: 3D shapes aligned by minimising the discrepancy between their centrelines, resulting in a tightly clustered set of vessels in which global anatomical pose is removed and only intrinsic morphological differences are retained.

### Correlation of shape and flow

#### Data and notation

For each subject *i* = 1*, …, n* (*n* = 12), we analysed *p* = 11 statistical shape model (SSM) coordinates **x***_i_* = (*x_i_*_1_*, …, x_ip_*)^⊤^ (latent modes) together with three wall shear stress (WSS) burden metrics 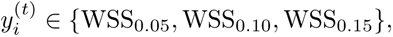 defined as the vessel surface area with WSS below the specified threshold (Pa). Each mode *x*_·_*_j_* was standardised across subjects (mean zero, unit sample standard deviation); WSS variables were analysed on their native scale. To localise associations within the shear distribution, we also formed disjoint bands B_0.05–0.10_ = WSS_0.10_ − WSS_0.05_ and B_0.10–0.15_ = WSS_0.15_ − WSS_0.10_.

#### Univariate rank-based associations

For each mode *j* and metric *t*, we computed *Spearman’s rank correlation ρ_S_* (*x*_·*j*_*, y*^(*t*)^), which is the Pearson correlation of mid-ranks and is robust to non-normality and monotonic nonlinearity.

#### Permutation testing (two-sided)

Given the small sample size (*n* = 12), we assessed significance by permutation. For each (*j, t*) we used the observed statistic *T*_obs_ = |*ρ_S_*| and generated a null distribution by permuting *y*^(^*^t^*^)^ across subjects while keeping *x*_·_*_j_* fixed (exchangeability under *H*_0_). With *B* = 2000 permutations, the *p*-value was

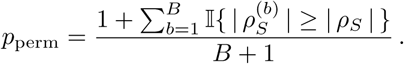

#### Multiple testing (BH–FDR)

We controlled the expected proportion of false discoveries across the family of mode × metric hypotheses using the *Benjamini–Hochberg* false-discovery rate (FDR) procedure applied to the permutation *p*-values. BH–FDR is appropriate under positive dependence, which is expected as WSS thresholds are nested and biological responses can involve multiple modes. We report FDR-adjusted *q*-values.

#### Uncertainty and stability

We computed 95% nonparametric bootstrap percentile confidence intervals for *ρ_S_*(resampling subjects with replacement; *B* = 2000). Robustness to single-subject influence was assessed by leave-one-out (LOO) re-estimation of *ρ_S_*; we report the LOO range 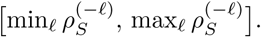

#### Scope of inference

Inference here is intentionally univariate. With *n* = 12 and *p* = 11, multivariate procedures (e.g., multivariate regression, MANOVA, CCA) are statistically unstable and prone to overfitting without substantial regularisation and external validation. Consequently, FDR-significant results identify the most prominent *individual* mode–metric associations; the absence of univariate significance for a mode should not be interpreted as absence of *joint* effect. Multivariate exploration is reserved for larger cohorts and is not used for inferential claims in this work.

#### Reporting

For each mode–metric pair we report: Spearman’s *ρ_S_*, permutation *p*_perm_, BH–FDR *q*-value, 95% bootstrap CI for *ρ_S_*, and the LOO range. Heatmaps display *ρ_S_* across modes and thresholds/bands, with cells flagged for *q <* 0.05.

## Results

This section presents research results following the structure of Materials and methods.

### Computational fluid dynamics

The analysis of haemodynamics focused on local variations due to shape differences and changes in boundary conditions. The metric of choice was the surface area of the wall of the vessel exposed to low wall shear stress (WSS) below three thresholds: 0.05, 0.10, 0.15 [*Pa*]. Low WSS has been associated with blood coagulation and clot formation [2] and is the assumed biophysical risk inference metric in this study.

Figure 6 provides an overview of the WSS calculated across the dataset. The top graph plots WSS area on the y-axis for each geometry on the x-axis. Three curves represent areas below different WSS cut-off values. Most cases have similar low-WSS areas, except *P* 7*_RCI_*, which shows a pronounced peak across all three thresholds. The area increases with the WSS threshold, giving vertically stacked curves - this is expected since the metrics are consecutive subsets of one another. Many geometries show sparse or patchy low-WSS regions. *P* 7*_RCI_* is prominent, with a notably larger low-WSS surface.

**Fig 6.**
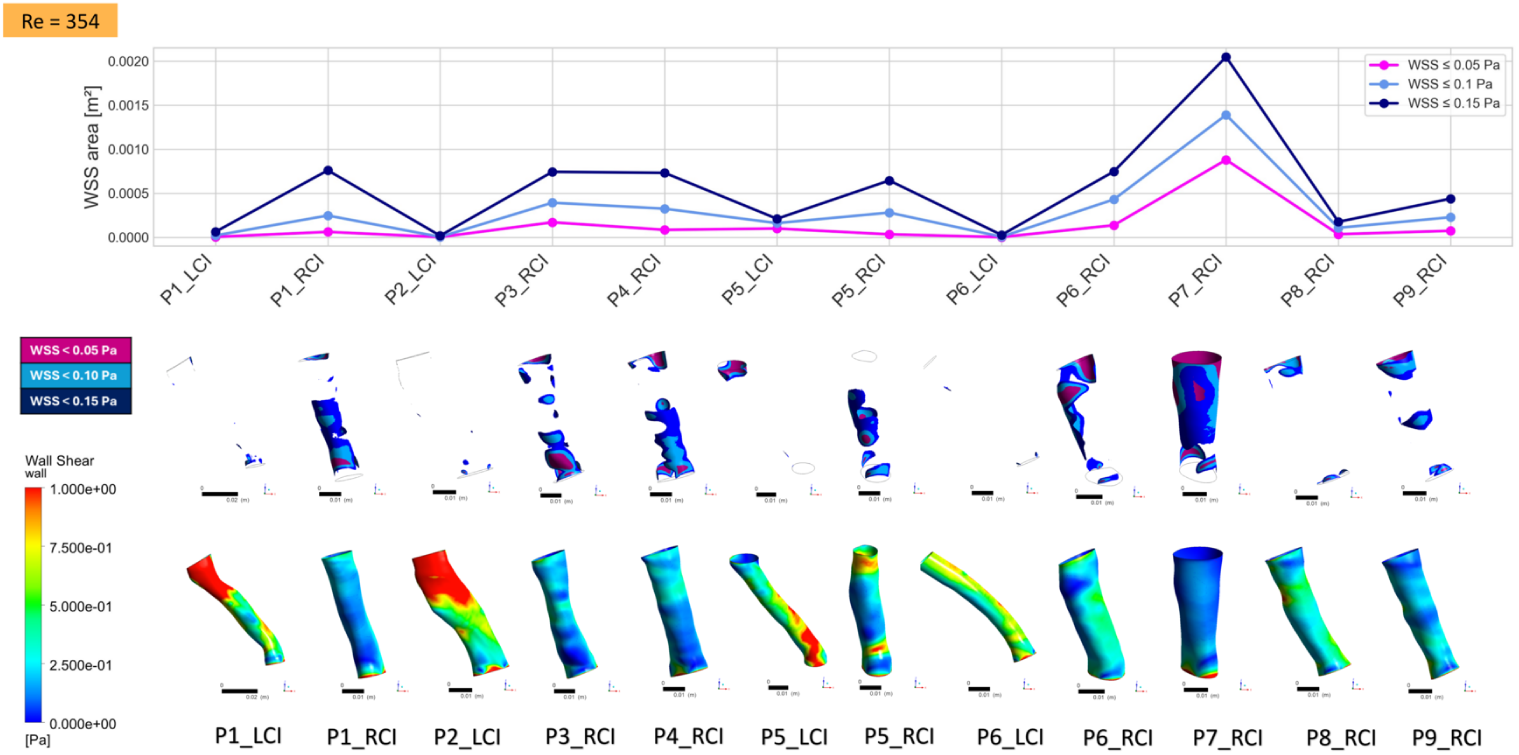
Steady-state wall shear stress metrics, computed for all shapes in the considered set; **bottom row:** WSS distribution between 0 and 1 Pa; **middle row:** WSS bands filtered by three thresholds (0.05, 0.10, 0.15 [Pa]); **top row:** surface area covered by WSS below the three thresholds. All simulations were performed with inlet velocity scaled to result in *Re* = 354.

In all cases, idealised geometries produced larger areas exposed to low wall shear stress compared to patient-specific geometries. On average, the difference between WSS area between 3D reconstructions and full 3D shapes was approximately 1.88 × 10^−4^ m^2^ for the threshold WSS ≤ 0.05 m^2^, 3.61 × 10^−4^ m^2^ for WSS ≤ 0.10 m^2^, and 3.86 × 10^−4^ m^2^ for WSS ≤ 0.15 m^2^. These values correspond to increases of approximately 118%, 136% and 78%, respectively, relative to the mean areas obtained from full 3D shapes. Although a small number of individual cases showed slightly smaller low-WSS regions in the 3D reconstruction (two to three cases depending on the threshold), the overall trend indicates that the simplified reconstruction systematically enlarges the extent of low-shear regions. The median differences followed the same pattern, ranging from 1.32 × 10^−4^ m^2^ to 3.50 × 10^−4^ m^2^ across the three thresholds, reinforcing the robustness of this effect. Representative WSS distributions for patient-specific and idealised 3D extruded geometries illustrate the systematic smoothing and redistribution of shear in the idealised models (see Supplementary material Fig. S3).

Figure 7 summarises the effect of ±20% inlet velocity perturbations on the patient-specific 3D models. Across all three thresholds, the absolute low–WSS areas increase with higher inflow and decrease with lower inflow. Importantly, the between-subject pattern is largely preserved: the same cases that exhibit the largest low–WSS area at baseline remain among the highest under both inflow perturbations, and subjects with small baseline area remain comparatively low. This rank stability indicates that, within the tested range, moderate inflow variations do not alter the relative haemodynamic risk ordering; rather, the geometric differences between subjects remain the primary driver of variability in low–WSS surface area.

**Fig 7.**
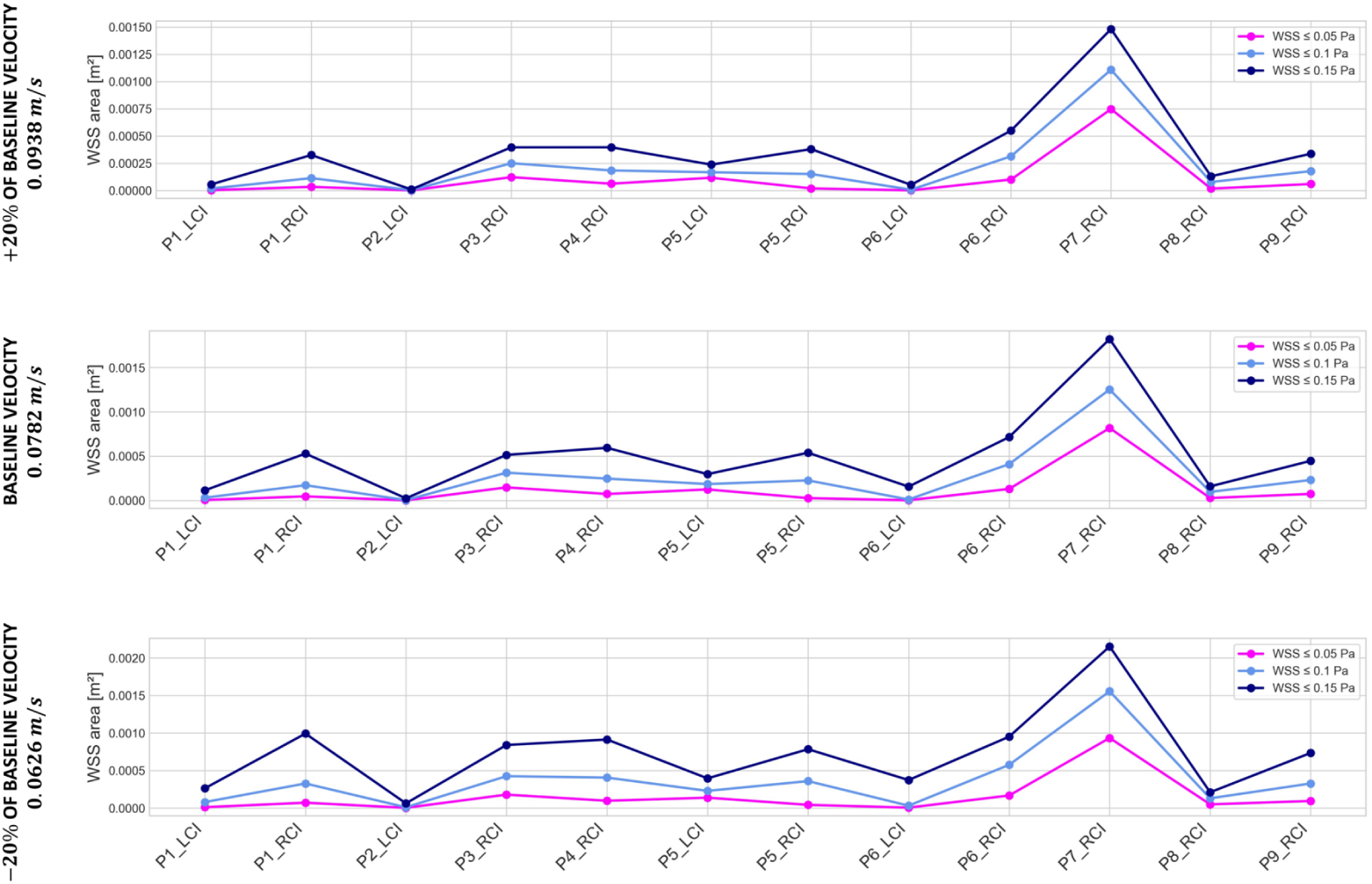
Sensitivity of low–WSS area to inlet velocity in patient-specific 3D geometries. Each panel reports the vessel wall area below three WSS thresholds (≤ 0.05*, ≤* 0.10*, ≤* 0.15 Pa) for all subjects, computed at 20% (bottom), baseline (middle), and +20% (top) of the baseline inlet velocity.

### Statistical shape modelling

The results of the shape analysis of 2D and 3D shapes are reported in dedicated subsections.

### 2D SSM

The set of 12 shapes considered was decomposed into 11 principal modes. Fig. 8 shows the first 5 modes (*m*1 − *m*5). The action of each mode is demonstrated by moving along the *i^th^* principal geodesic mode by *k* standard deviations *σ*, with 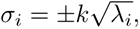 where *k* is an integer and *λ* is the corresponding eigenvalue. Increasing *k* produces statistically significant variations along a principal mode. In the case of the blood vessels considered, ±2*σ* are assumed to be anatomically plausible and therefore further used for CFD simulations.

**Fig 8.**
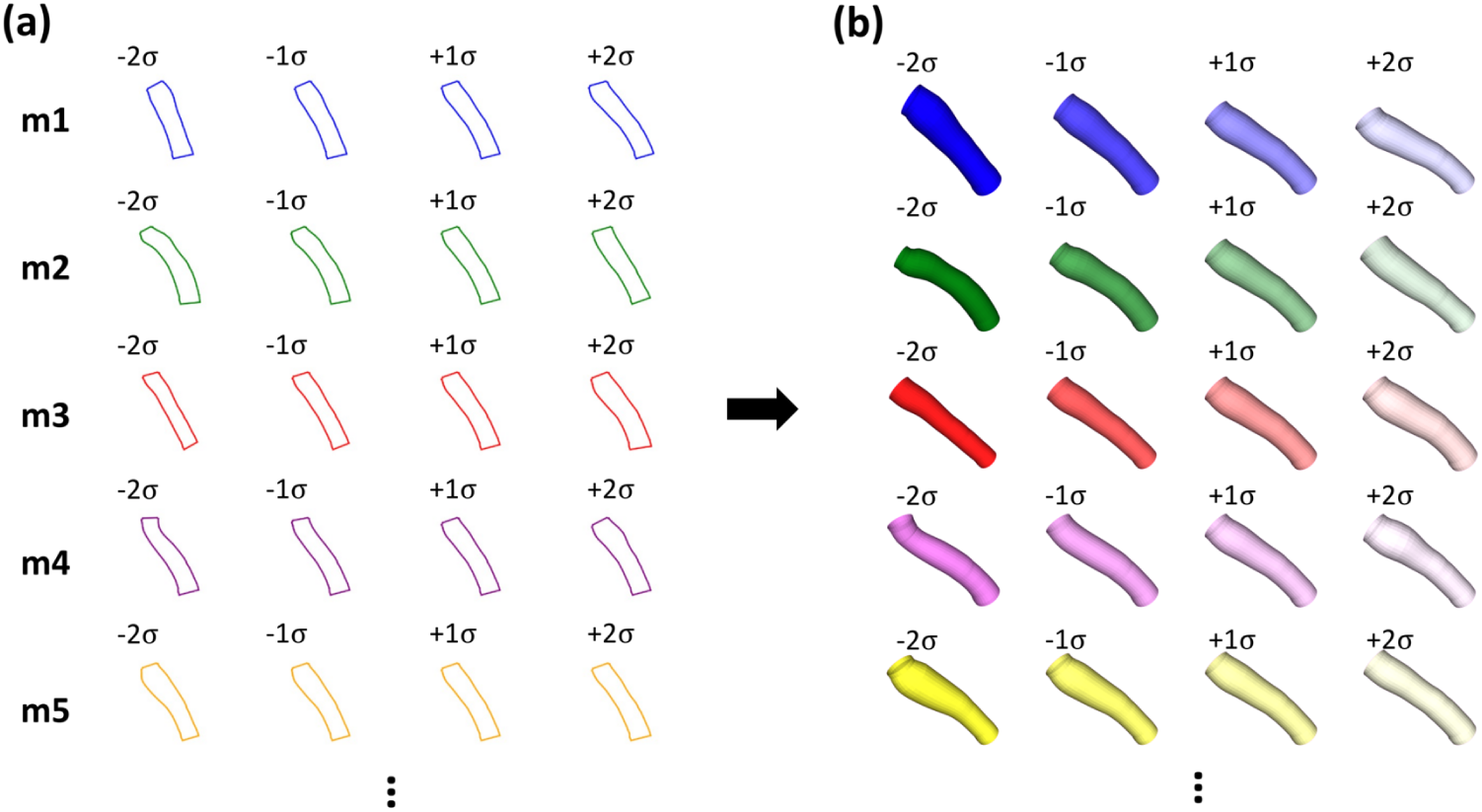
(a) A subset of shape modes from the 2D SSM analysis with a subset of shapes obtained by moving along each principal geodesic mode by a certain number of standard deviations, *σ*, from the mean shape. (b) Corresponding 3D extrusions, as used in our CFD analysis.

The action of the first 5 modes may be described as follows:

- **Mode 1:** A combination of global bending and subtle changes in calibre; as the deformation moves from −2*σ* to +2*σ*, the vessel becomes straighter while exhibiting mild co-varying bulging and thinning along its length.
- **Mode 2:** Primarily governs proximal–distal tapering; negative values produce a bulkier, more uniformly thick tube, whereas positive values create a more tapered and slender geometry, with little effect on overall curvature.
- **Mode 3:** Captures localised irregularity versus smoothing; at −2*σ* the midsection appears thicker or more swollen, while at +2*σ* the shape becomes smoother and more uniformly cylindrical, with curvature largely preserved.
- **Mode 4:** Represents a mixture of secondary bending or twisting and local bulging; negative excursions introduce an accentuated S-shaped curvature together with a noticeable swelling, while positive excursions reduce both curvature and bulge.
- **Mode 5:** Modulates the distal calibre; movement from −2*σ* to +2*σ* narrows and then widens the distal end, producing a funnel-like form, with limited influence on global bending.

The explained variance calculated for each mode revealed that *m*1 and *m*2 explain 33% and 21% of the variance respectively. The remaining modes describe between 3% and 7% each (Fig. 9).

**Fig 9.**
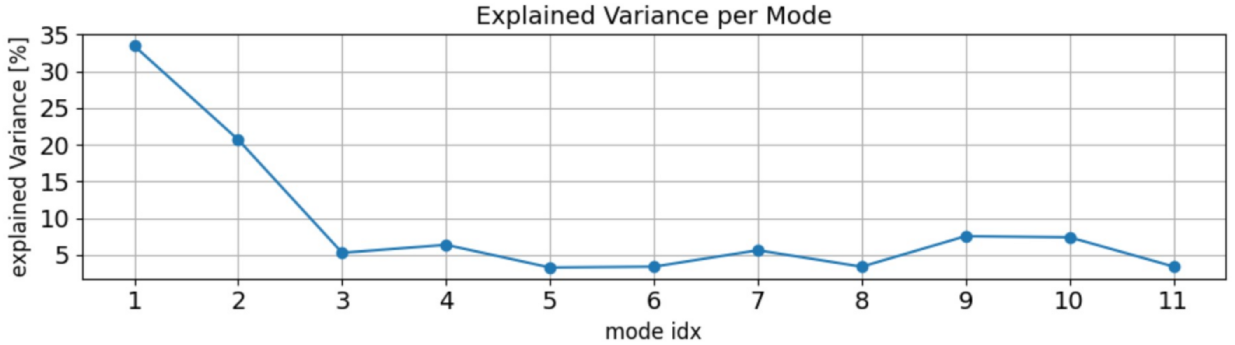
Explained variance per shape mode for SSM applied to 2D shapes.

### 3D SSM

The deterministic atlas step produced a population template (Fig. 10) that reflects the average 3D iliac vein morphology in all subjects. This mean geometry forms the baseline from which each principal geodesic mode is derived, allowing the subsequent analysis to characterise how individual shapes deviate from the cohort mean.

**Fig 10.**
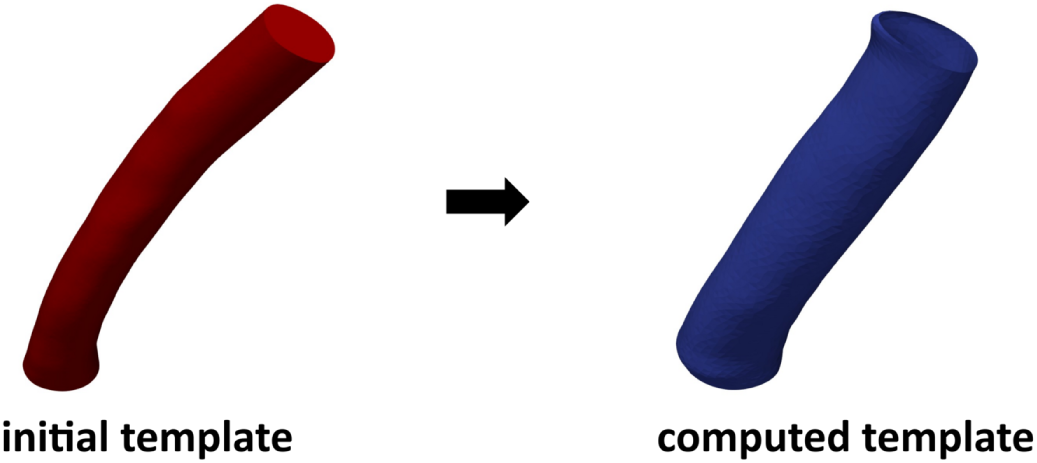
Initial template geometry - one of the subjects (left) and template obtained from the deterministic atlas step (right) for the 3D iliac vein cohort. This shape represents the mean anatomy and serves as the reference for Principal Geodesic Analysis.

Following centreline-based alignment, the venous geometries exhibited very limited inter-subject variability, with the shapes overlapping closely and showing no substantial global deformations. The principal geodesic analysis therefore revealed subtle morphological differences. The first mode, which explained 17.4% of the total variance, corresponded to a largely uniform change in lumen calibre and reflected the primary source of anatomical variability within the cohort. All subsequent modes displayed a remarkably flat variance spectrum (each accounting for approximately 7–9%), indicating the absence of any dominant secondary deformation pattern. Modes 2 and 3 primarily described slight adjustments in axial taper and very mild residual curvature, whereas Modes 4–8 captured localised radial variations, including small mid-segment expansions, weak axial shifts of these expansions, and minimal deviations from circularity. Higher-order modes (Modes 9–11) represented distributed micro-scale geometric effects, such as subtle modifications of surface curvature and weak orthogonal bending or twist-like deformations. Collectively, these modes characterise fine-scale, spatially diffuse shape variability that remains after stringent alignment, consistent with the smooth tubular anatomy of the deep veins. Fig. 11 shows the distribution of explained variance across shape modes.

**Fig 11.**
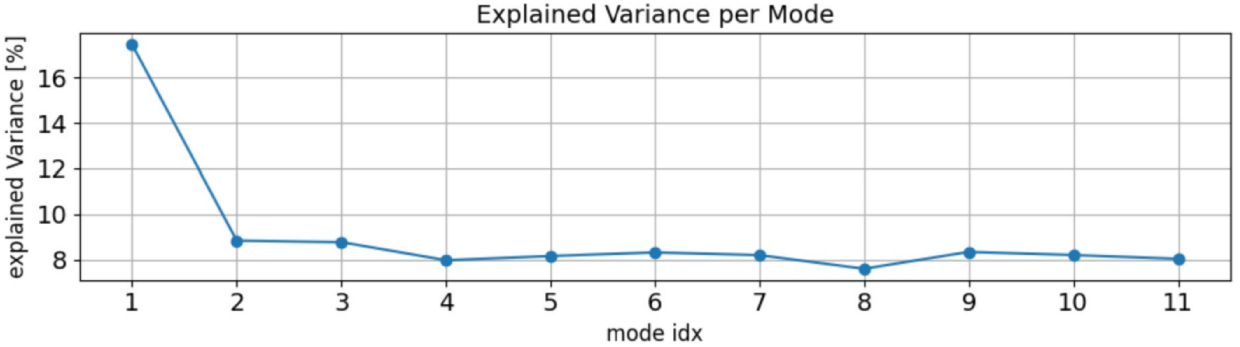
Explained variance per shape mode for SSM applied to 3D shapes.

To visualise the 3D effect of the dominant principal geodesic, Fig. 12 colours the template surface by the surface-normal displacement magnitude induced at −2*σ* and +2*σ* along Mode 1. The response is spatially smooth and of small amplitude (maximal |Δ*n̂*| in the sub-millimetre range), consistent with the strong centreline-based alignment that removes global pose and curvature. The displacement hotspots track gentle calibre/taper adjustments along the vessel, with near-symmetric patterns at −2*σ* and +2*σ*, indicating that Mode 1 acts primarily as a distributed dilation–constriction rather than introducing sharp local bulges.

**Fig 12.**
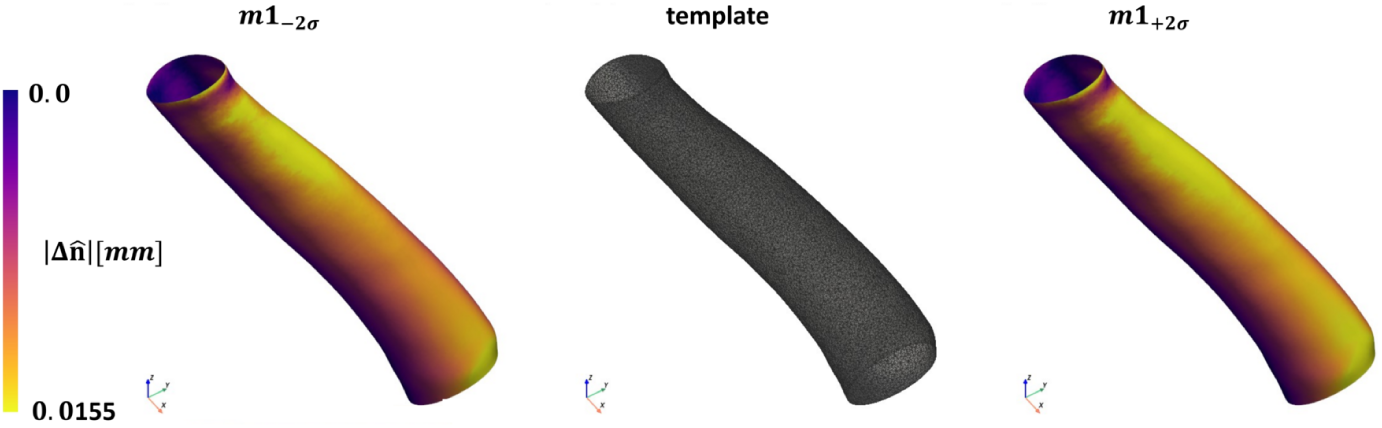
Three-dimensional morphological response of the template to the first principal geodesic mode (m1). Left and right panels: magnitude of surface-normal displacement Δ*n̂* (mm) relative to the template when moving to –2*σ* and +2*σ* along m1, shown with a shared colour scale. Middle: the atlas template (mean) geometry for reference. The deformations are spatially smooth and sub-millimetric, indicating that m1 primarily modulates local lumen calibre/taper, consistent with the centreline-based alignment.

### 2D SSM vs 3D SSM

A clear distinction emerges between the 2D and 3D statistical shape models (SSMs), primarily due to differences in the underlying alignment procedures and the geometric information available in each representation. In the 2D SSM, the Procrustes-type alignment preserved a substantial degree of global pose variation, as evidenced by the spread of the outlines, resulting in the first two modes capturing large-scale effects such as bending, tapering and co-varying calibre changes. Consequently, Modes 1 and 2 explained 33% and 21% of the total variance, respectively, with subsequent modes representing increasingly localised features. In contrast, the 3D SSM employed a centreline-based alignment that effectively removed all global pose and orientation differences, producing a tightly clustered set of shapes with minimal residual deformation. As a result, the 3D modes primarily describe subtle, spatially diffuse variations in local radius, taper and surface curvature, with only the first mode exhibiting a modest dominance (17.4% of variance) and the remaining modes displaying an unusually flat spectrum (approximately 7–9% each). Thus, while the 2D modes reflect both global and local geometric variability, the 3D modes isolate fine-scale anatomical nuances that remain after stringent alignment, consistent with the smoother, more tubular nature of the underlying venous anatomy.

### Correlation of shape and flow

The results of 2D SSM and 3D SSM are compared and contrasted in dedicated subsections.

### SSM of 2D projections of the veins

Fig. 13 shows the distribution of WSS computed in simulations for a subset of shape modes. Across all three WSS thresholds, modes 4, 7, and 8 are the most sensitive to changes in wall shear stress (WSS), capturing medium-scale geometric features such as local curvature or diameter modulation. In contrast, modes 6 and 9 show minimal changes across the range of 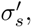 indicating low sensitivity. Although the trends are not strictly linear or monotonic, consistent behaviours emerge: mode 4 progressively expands the low-WSS regions from −2*σ* to +2*σ*, while mode 7 shows a steady reduction of these regions over the same range. Mode 8 produces the largest overall decrease in low-WSS area, particularly for the threshold *<* 0.15 Pa. Mode 1 shows moderate sensitivity; it consistently decreases the low-WSS area from −2*σ* to +2*σ* and tends to produce large mean WSS-affected regions, yet it also generates the greatest variation in the shape of thresholded bands, consistent with its role in the description of large-scale global deformations. Across all thresholds, the −2*σ* geometries induce the strongest CFD effects, likely because these shapes are more extreme—with pronounced bulging, narrowing, or bending near vessel outlets—leading to stronger sensitivity of WSS to local curvature and diameter perturbations.

**Fig 13.**
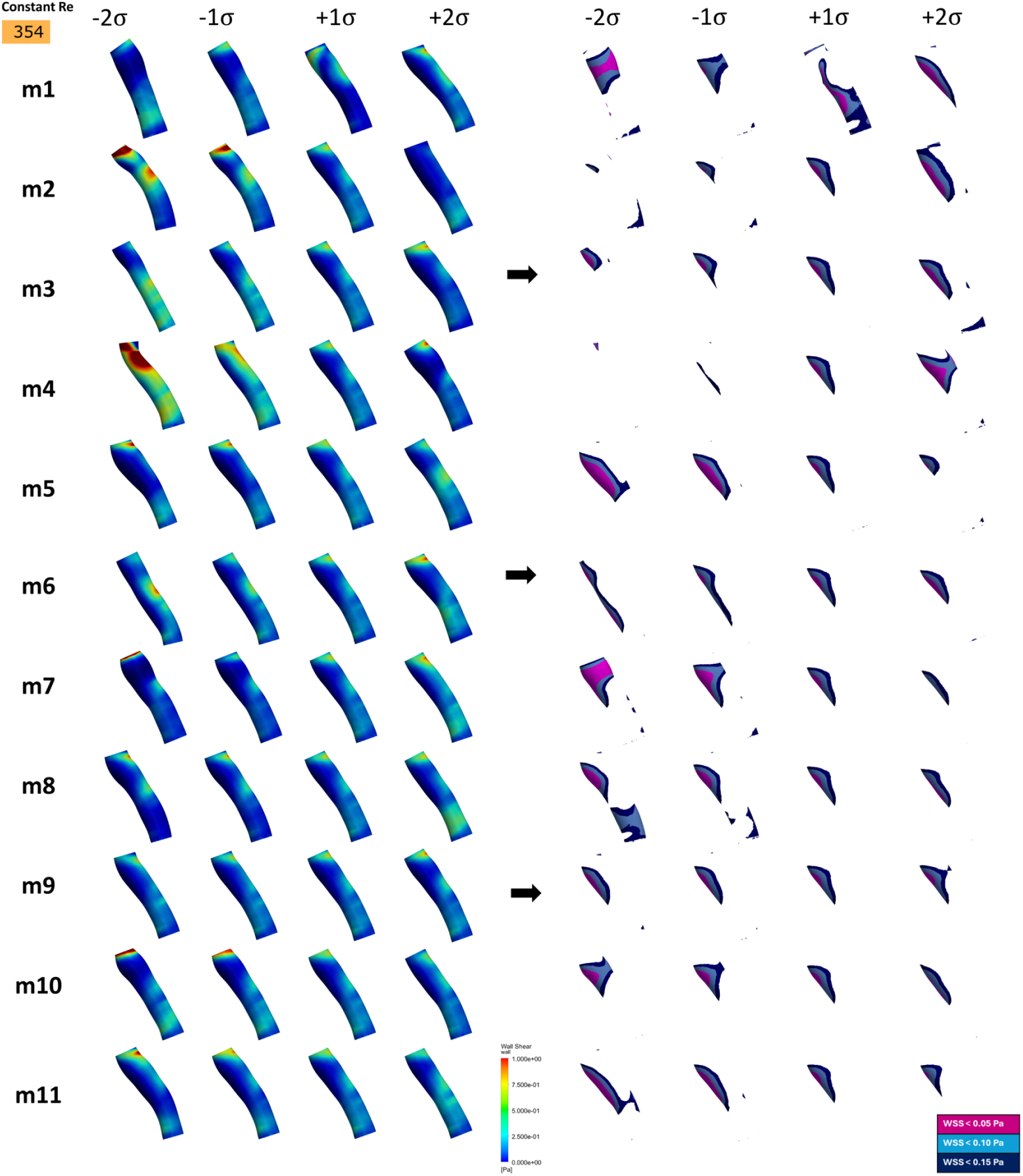
WSS distributions for shapes varied along a set of shape modes, *m_i_*.

Fig. 14 summarises the *σ*–response of the low–WSS burden within the 2D SSM. Across all thresholds, Mode 1 exhibits the largest dynamic range, with clear changes in the low–WSS area as *σ* varies, while most other modes show smaller, more muted responses.

**Fig 14.**
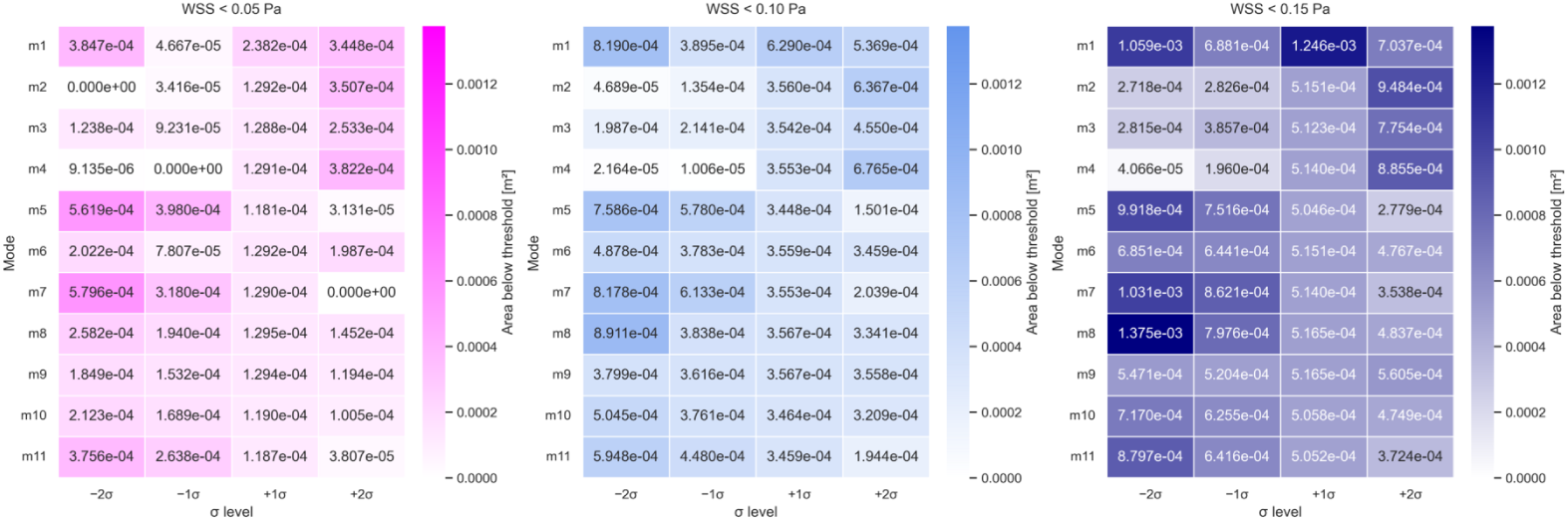
Mode–response of low–WSS burden in the 2D SSM. Each heatmap reports the vessel wall area below a given WSS threshold (left: ≤ 0.05 Pa; middle: ≤ 0.10 Pa; right: ≤ 0.15 Pa) for shapes obtained by moving along each principal mode by *σ –*2*, –*1, +1, +2, relative to the 2D template. Rows correspond to modes (m1,…, m11); columns correspond to the *σ* level. Stronger cell-to-cell contrast indicates greater haemodynamic sensitivity to movement along that mode. Colour scales are kept per-threshold and shared across all modes.

Fig. 15 complements these findings by positioning the *σ* offset shapes in selected latent subspaces (formed by the first three most-correlated modes for each metric) and colouring the points by the associated WSS area value.

**Fig 15.**
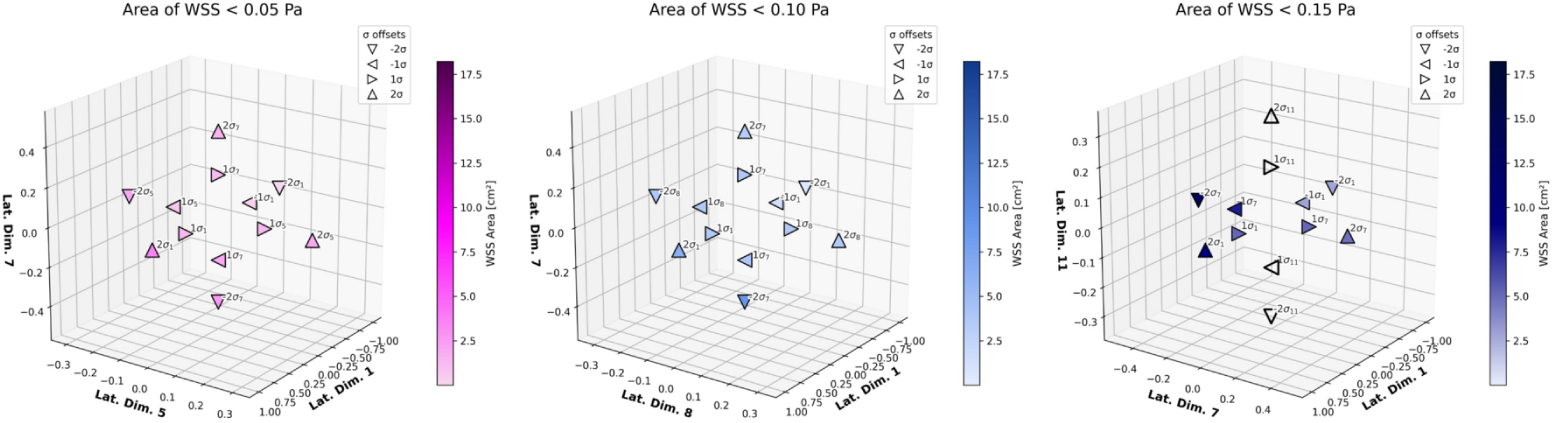
Low–WSS across *σ*–displacements in selected 2D SSM subspaces. Each panel shows the positions of the *σ*–offset shapes (–2*σ*, –1*σ*, +1*σ*, +2*σ*; marker shapes) in a 3D plot of the 2D latent coordinates (three chosen modes per panel), with marker colour denoting the corresponding low–WSS area (m^2^) at the indicated threshold.

### SSM of 3D shapes

Fig. 16 illustrates the thresholded WSS as the geometry is moved along Mode 1. Relative to the template, *m*1_−2_*_σ_* reduces the low-WSS area across all thresholds (0.05 Pa: −38%; 0.10 Pa: −27%; 0.15 Pa: −44%). In contrast, *m*1_+2_*_σ_* leaves the *<* 0.05 Pa band essentially unchanged and shows only a small decrease at *<* 0.10 Pa (−3%), but produces a substantial increase at *<* 0.15 Pa (+73%). The largest swing between the extremes occurs for the *<* 0.15 Pa band (+212% from *m*1_−2_*_σ_* to *m*1_+2_*_σ_*), indicating that m1 predominantly expands moderately low-shear regions rather than generating additional very-low WSS patches. This threshold-dependent behaviour is consistent with the smooth, sub-millimetric calibre/taper changes captured by Mode 1 in the 3D displacement map (Fig. 16).

**Fig 16.**
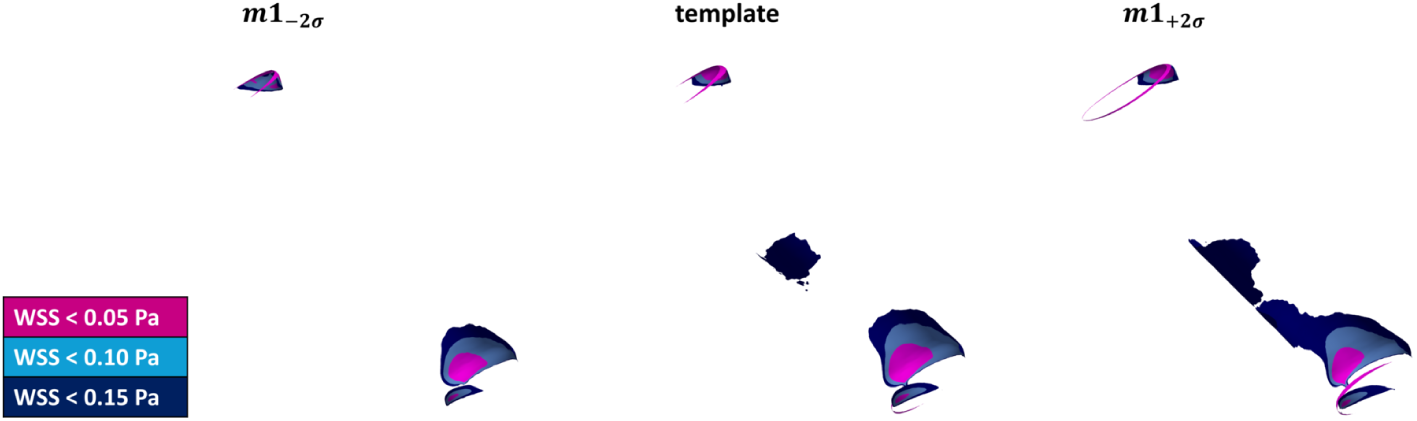
Effect of the first 3D shape mode (m1) on the distribution of low wall shear stress (WSS) over the venous surface. The geometry is shown at *m*1_−2_*_σ_* (left), the atlas template (centre), and *m*1_+2_*_σ_* (right).

### Comparison of correlations between SSM and CFD

Fig. 17 reports the association between SSM modes and CFD metrics from both the 2D and 3D SSM analysis. In the 2D case, Mode 1 shows consistently strong negative correlations with all WSS metrics (*ρ_Spearman_* ranging from –0.58 to –0.93), and remains statistically significant for the 0.10 Pa and 0.15 Pa thresholds and their corresponding WSS bandwidths after permutation testing with Benjamini–Hochberg FDR correction. This indicates that the dominant direction of variation in the 2D latent space captures geometric features that are strongly associated with low-WSS regions.

**Fig 17.**
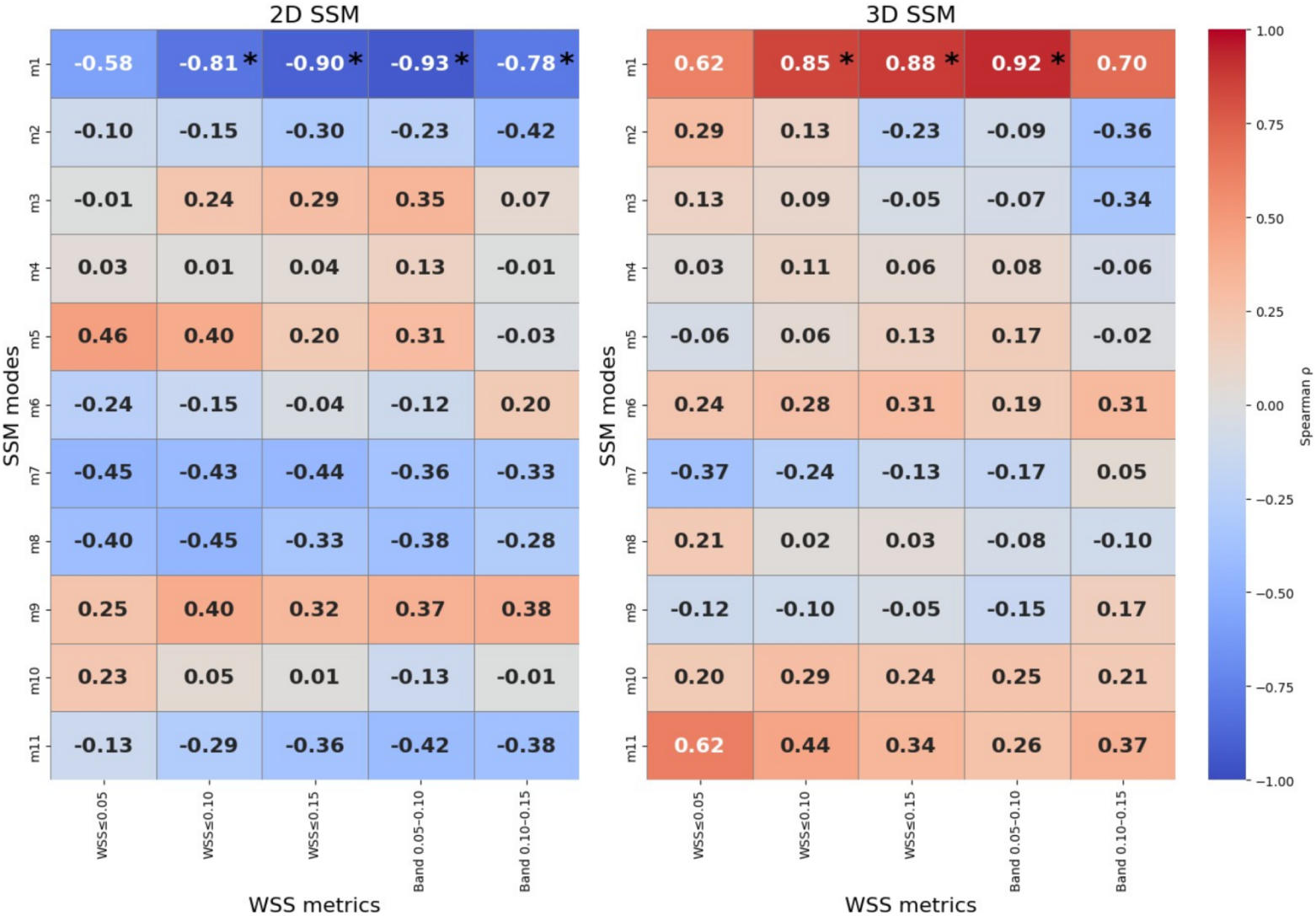
Spearman rank correlations between statistical shape model (SSM) modes and CFD-derived wall shear stress (WSS) metrics for the 2D and 3D models. Each heatmap shows the correlation coefficient (*ρ_Spearman_*) between individual SSM modes (rows) and WSS burden metrics (columns), including absolute thresholds (*WSS* ≤0.05, *WSS* ≤0.10, *WSS* ≤0.15) and derived WSS bandwidths (Band 0.05–0.10, Band 0.10–0.15). Asterisks mark associations that remain significant after permutation testing with Benjamini–Hochberg false-discovery-rate correction (q < 0.05). The 2D SSM displays a single dominant flow-associated direction (Mode 1), whereas the 3D SSM exhibits weaker and more heterogeneous associations distributed across multiple modes.

In contrast, the 3D SSM demonstrates a markedly different pattern: correlations with individual shape modes are weaker, more variable in sign, and distributed across several modes, with no association surviving FDR correction. This diffuse correlation is consistent with the substantially flatter variance spectrum of the 3D model, where anatomical variability is spread across many modes rather than concentrated in one or two dominant directions. As a result, any flow-related differences are distributed more broadly across the 3D latent space, reducing the strength of univariate associations in this sample.

### Visualisation of SSM and CFD correlations

Fig. 18 shows the variation of low-WSS regions with two selected shape modes from the 2D SSM analysis. In this figure the left column uses the shape modes which contribute most to the overall shape variance (Fig. 9) and the right column uses the shape modes with the highest Spearman rank correlation (Fig. 17). The left column shows that most subjects cluster near the origin with a small number of cases occupying the extremes of Mode 1. These extremes coincide with the largest low–WSS areas (taller bars), indicating that the dominant 2D direction captures a substantial share of the geometry that co-varies with WSS. By contrast, the right column yields a subspace where separation between high- and low-burden cases is more apparent: subjects with the greatest low-WSS areas tend to sit at one corner of the plane, while low-burden subjects sit at the opposite corner. This pattern is consistent across thresholds (≤ 0.05, 0.10, 0.15 [Pa]), with the contrast most visible at 0.15 Pa where between-subject spread in bar height is greatest. Taken together, the figure illustrates that (i) a projection onto the first two principal 2D modes is already informative for WSS ordering, and (ii) selecting modes by *Spearman correlation* with each WSS threshold further sharpens the visual separation of subjects along directions most relevant to that threshold.

**Fig 18.**
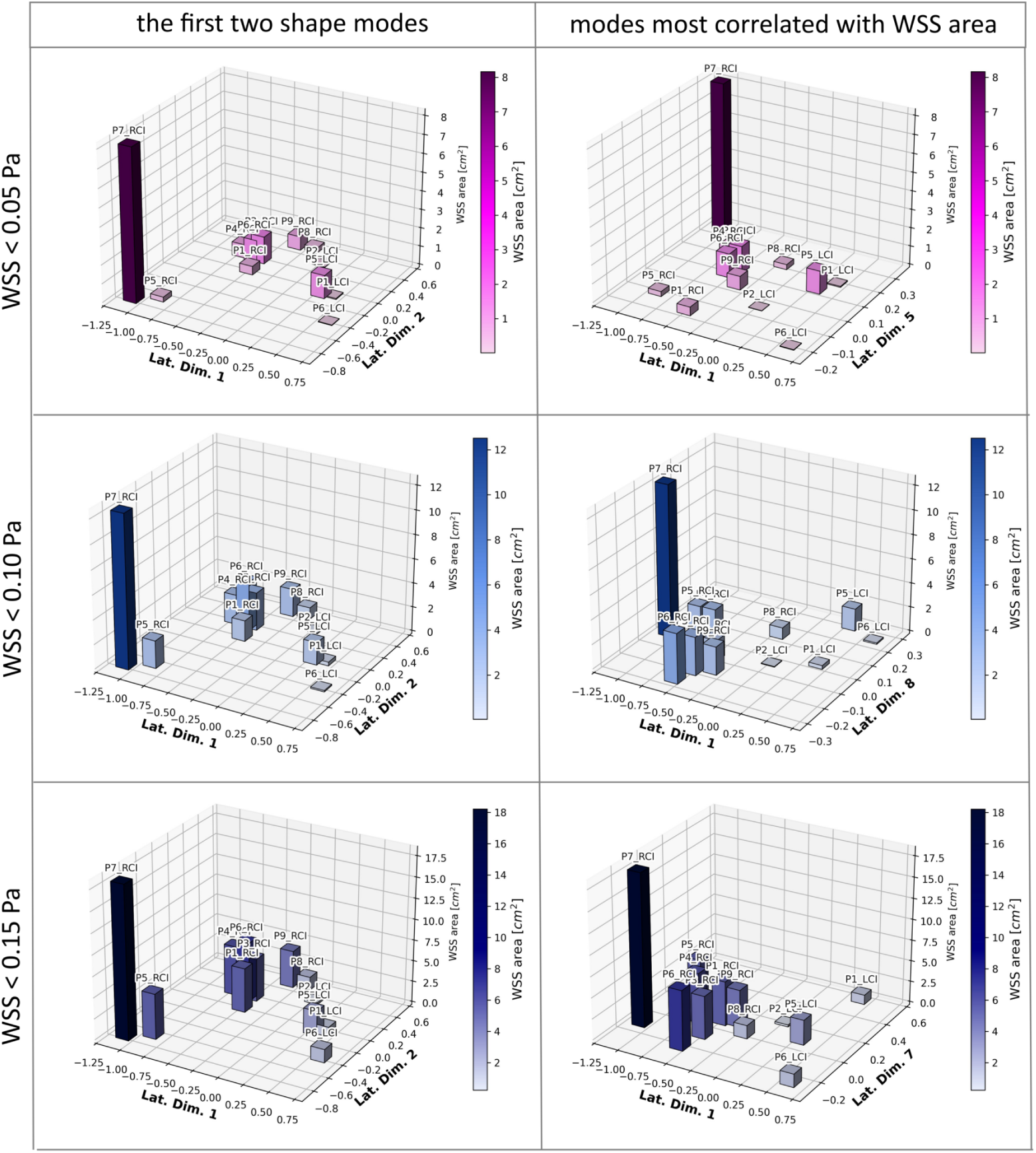
Distribution of shapes in a 2D subspace defined by two shape modes - according to SSM performed on 2D shapes (left column) and most correlated with CFD outputs (right column) - for three WSS area thresholds.

## Discussion

This study reports the combination of SSM methods and CFD analysis, to evaluate the variation in regions of low-WSS with venous anatomy and the sensitivity of these associations to both the form of the medical images used and the processing of the images during shape analysis.

The Deterministic Atlas enables the construction of a population-based template and the alignment of individual geometries to a common reference frame. This ensures that subsequent PGA (Principal Geodesic Analysis) captures genuine morphological variation rather than differences arising from scale, orientation, or arbitrary positioning. The atlas encodes inter-subject deformations into a structured parameter space, allowing PGA to identify dominant modes of variation in a mathematically consistent manner. This provides a compact representation of population-level shape differences that can subsequently be examined in relation to additional modelling outputs.

CFD simulations provide complementary physics-based estimates of how these variations influence local flow patterns and the distribution of wall shear stress. Because CFD is performed independently of the SSM construction and under consistent boundary conditions, it isolates the effect of geometry from that of physiological or inflow-related factors. Integrating statistical shape analysis with physics-based modelling therefore enables a mechanistic interpretation of the geometric modes identified by PGA and facilitates a more comprehensive assessment of how anatomical variability may relate to metrics that are relevant for thrombosis research.

### Shape variance

Although both the 2D and the 3D statistical shape models (SSMs) were constructed from the same set of venous anatomies, the latent spaces they produce differ substantially. Two methodological factors explain this divergence: (i) the variance structure inherent to each model, and (ii) the way shapes were aligned prior to SSM construction. The 2D SSM exhibited a strongly hierarchical decomposition of geometric variability, with Mode 1 explaining approximately 33% of the total variance and Mode 2 around 20%, while all remaining modes contributed between 3–7% each. In contrast, the 3D SSM distributed its variance much more evenly: Mode 1 explained only 17%, and all other modes contributed roughly 8% each. This contrast is important: in the 2D model, a large portion of anatomical information is compressed into the first two modes, whereas in the 3D model, geometric differences are spread across a relatively flat latent spectrum (Figures 9, 11).

### Impact of shape alignment

These structural differences are further amplified by the fundamentally different alignment procedures applied to the 2D and 3D shape sets. An alignment preserving anatomical orientation was possible in our case because the data originated from MRI/CT acquisitions, allowing us to register shapes in a way that respects their *in–situ* relationships. This anatomical alignment was preserved in the case of the 2D SSM, making the resulting latent space more anatomically interpretable. By contrast, if the 2D geometries had been extracted solely from angiography (particularly when only limited or variable frame geometry is available), there would be far less inherent information about absolute orientation and pose. Under such circumstances, anchoring shapes to a fixed anatomical frame might not be possible, and a more abstract alignment strategy would be required.

For the 3D SSM, an anatomically anchored alignment would, in principle, preserve the natural inter–subject anatomical spread. However, with our current data and tooling, we were unable to identify a set of SSM parameters that led to stable convergence under such an alignment in Deformetrica. Consequently, we employed a centreline–based alignment, which minimises inter–subject differences by bringing venous centrelines into close correspondence, effectively placing the veins “in a vacuum”. This has two main consequences: first, it reduces positional variability between subjects, aiding numerical stability and comparability; and second, it suppresses a portion of natural anatomical dispersion. Clinically, centreline–based normalisation is arguably more robust across diverse imaging contexts, as it aligns shapes within the vessel’s intrinsic coordinate system rather than relying on a potentially inconsistent external frame.

In short, anatomical alignment is ideal in theory because it preserves native anatomical variation and orientation; however, centreline alignment is often more pragmatic, particularly in 3D, where algorithmic stability and cross–cohort robustness matter. In 2D, both alignment philosophies can be explored (and MRI–based data make anatomically anchored registration feasible). In 3D, with our present data and software, only the centreline–based approach converged reliably. We acknowledge that this means that the 2D and 3D SSMs were built under different alignment regimes, but this reflects the practical constraints of the imaging sources and current modelling capabilities.

### Consequences for shape–flow relationships

Using the original WSS fields for both models, the 2D and 3D SSMs produced different sets of modes that correlate with haemodynamic indicators, and these differences align with the manner in which each model encodes geometric variation. Because the three WSS thresholds are cumulative, nested, and therefore strongly correlated, each threshold should not be interpreted as an independent outcome but rather as a different cut-off applied to the same underlying distribution. Accordingly, a single Benjamini–Hochberg (BH) false-discovery-rate (FDR) correction was applied across all mode–threshold tests.

In the 2D SSM, a single geometric direction—Mode 1—emerged as the dominant flow–associated mode. It showed strong correlations with low-WSS area across the considered thresholds, reaching the highest Spearman coefficients (*ρ*_S_ ≈ 0.85–0.88) and surviving permutation-based significance testing and FDR correction for the 0.10 Pa and 0.15 Pa thresholds. Importantly, the use of permutation testing ensured exact small-sample inference given *n* = 12, and the FDR procedure identified Mode 1 as the most prominent *individual* association among the family of tests. Although other modes appeared among the top correlations, their *q*-values did not reach the FDR threshold; however, this does *not* imply that these remaining modes have no joint or subtle influence on WSS metrics. Rather, it reflects that, at this sample size, their individual effects cannot be distinguished from chance with high confidence.

Conversely, in the 3D SSM, which has a much flatter variance spectrum, no single dominant shape–flow direction was isolated. Correlations between 3D latent coordinates and WSS were weaker, more heterogeneous across modes, and none survived FDR correction. This dispersion is consistent with the underlying shape variance being spread more evenly across modes: if geometry is decomposed into many comparably weighted directions, any flow-related variation is likewise distributed, making it less likely for any single univariate association to surpass statistical significance in a small sample.

Together, these findings show that the 2D SSM distilled a single, dominant geometric direction that strongly co-varied with the WSS-derived metrics, whereas the 3D SSM captured a broader and more diffuse set of shape differences, each accounting for a smaller fraction of the anatomical variation and showing weaker or inconsistent associations with WSS. These differences originate in the representation of shape variability in the latent spaces, rather than from any differences in the CFD fields themselves. The univariate statistical findings should therefore be interpreted as highlighting the most prominent single-mode associations rather than implying the absence of more complex, multivariate relationships.

### The impact of sample size

The current dataset of 12 shapes limits robust statistical inference when studying subtle, multi–dimensional relationships between shape and flow. With this anatomical dataset more nuanced effects may not be detectable and generalisation to broader populations requires further work. Increasing the sample size would allow more stable and reliable correlation estimates, narrower confidence intervals, improved detection of moderate effects, more reliable mode ranking, and more meaningful multivariate modelling. Greater numbers would also reduce variance in both the shape space and the CFD-derived metrics.

### Clinical and mechanistic meaning of shape-based prediction

While the present study demonstrates that certain geometric features can be associated with variation in WSS metrics, translating such associations into clinically actionable prediction requires larger cohorts, validation against independent datasets, and ideally the inclusion of longitudinal information. The current results should therefore be interpreted as an exploratory assessment of how anatomical variability manifests in CFD-derived quantities, rather than as a definitive shape-based risk model.

### Summary of limitations

This study has several limitations. First, the sample size (*n* = 12) constrains statistical power with weaker effects unlikely to be detected, uncertainty intervals wide and rankings of shape modes which may be unstable under resampling. Inference was therefore restricted to univariate, permutation-tested Spearman correlations with BH–FDR control; multivariate inference was not attempted, as *n* ≈ *p* renders such procedures statistically fragile and prone to overfitting without strong regularisation and external validation. While the sample size constrains statistical power, our use of permutation tests, bootstrap confidence intervals, and leave-one-out robustness analysis ensures that the reported associations represent the strongest detectable geometric-haemodynamic relationships under the available data. This study uses a sample size significantly greater than previous studies of patient-specific venous haemodynamics in the iliac veins REF.

Secondly, the three WSS thresholds are nested and highly correlated; while this justifies a single FDR family across mode-by-threshold tests, it also means that threshold-specific findings should be interpreted as different windows on the same underlying distribution rather than independent outcomes. Thirdly, the 2D and 3D SSMs were built under different alignment regimes (anatomically anchored in 2D; centreline-based in 3D), reflecting the data sources and algorithmic convergence constraints. Consequently, their latent spaces are not directly comparable.

Finally, CFD boundary conditions were held consistent to isolate geometry, but residual sensitivities (e.g., meshing, solver tolerances) cannot be fully excluded. Uniform outlet pressure simplifies collateral return and was chosen to standardise comparisons; consequently, results reflect geometry-driven differences rather than patient-specific haemodynamics. Overall, results should be viewed as exploratory and hypothesis-generating, motivating larger cohorts, harmonised alignment strategies where feasible, and confirmatory multivariate analyses with a larger sample size.

## Conclusion

This work establishes a mechanistic link between venous morphology and haemodynamics by integrating statistical shape modelling with CFD across two levels of geometric fidelity. We show that fidelity and alignment choices fundamentally reshape the latent structure of anatomical variability and, in turn, the apparent strength and localisation of shape–flow relationships. In 3D, the dominant mode produces smooth, submillimeter calibre/taper adjustments that measurably modulate low WSS regions, broadening moderately low-shear regions without generating new very-low WSS pockets; these effects persist under moderate inflow perturbations, indicating geometry—rather than boundary-condition choice within a physiological range—as the primary driver of haemodynamics. Together with the systematic inflation of the low-WSS area in simplified geometries, our results caution that model selection (projection, extrusion, or full 3D; and alignment strategy) can qualitatively alter haemodynamic inferences. This work represents the first detailed haemodynamic examination of patient-specific flow in the iliac veins, describing a reproducible and general computational workflow which maps principal deformations to localised regions of low WSS. Future work will scale the cohort, incorporate and extend to multivariate or surrogate modelling to derive robust, clinically useful shape biomarkers of haemodynamic risk in DVT.

## Acknowledgments

This work was supported by the National Science Centre (NCN) under Preludium’22 grant agreement UMO-2023/49/N/ST6/04252.

This project has received funding from the European Union’s Horizon 2020 research and innovation programme under grant agreement No 857533 and was created within the project of the Minister of Science and Higher Education “Support for the activity of Centers of Excellence established in Poland under Horizon 2020” on the basis of the contract number MEiN/2023/DIR/3796 and is supported by Sano project carried out within the International Research Agendas programme of the Foundation for Polish Science, co-financed by the European Union under the European Regional Development Fund.

## Data availability

Dataset associated with this publication available at https://doi.org/doi:10.71580/SANO/GVPFQ5.

